# APV-Sankey: A Comprehensive Toolbox for Aptamer Screening and Visualization

**DOI:** 10.1101/2025.02.11.637585

**Authors:** Yu Zhang, Yashan Wang, Yajing Gao, Keyi Hu, Hao Gong, Haijing Jia, Xin Zhang, Xinhui Lou

## Abstract

Aptamers, short single-stranded DNA or RNA molecules, have gained prominence as molecular recognition elements in diagnostics and therapeutics. Screening high-performance aptamers from SELEX process is tough due to limited library diversity, PCR bias, and low library enrichment efficiency. The enriched highest frequency sequences often do not have the highest affinity or specificity. Thus, we developed APV-Sankey, a novel and versatile toolbox designed for the rapid aptamer screening and visualization of the enrichment process. Its key feature is the innovative use of Sankey charts for interactive and informative visualizations. These charts facilitate tracing the evolution of K-mers (K-mer is just a sequence of k characters in a string) across SELEX rounds, aiding in the identification and selection of high-affinity sequences containing K-mers while excluding high-frequency sequences without affinity. Besides integrating essential functions for aptamer analysis, we also developed advanced methods including K-mer concatenation and K-mer evolution, combined with Sankey chart visualizations. Using this toolbox to analyze experimentally screened sequences, we successfully identified high-performance candidate aptamers of rapamycin and thrombin.

## Introduction

Aptamers are short, single-stranded oligonucleotide sequences, typically ranging from 30 to 100 nucleotides in length [1]. In the rapid development of molecular biology and biotechnology, aptamers have become recognized as versatile molecular recognition elements with applications ranging from diagnostics to targeted therapeutics [2–4]. Typically, high-performance aptamers are developed using an in vitro selection technique known as Systematic Evolution of Ligands by Exponential Enrichment (SELEX) [5, 6]. Through various separation methods, sequences with high binding affinity are selected and amplified via Polymerase Chain Reaction (PCR), followed by the preparation of single-stranded secondary library. This cycle is repeated multiple times to enrich the candidate sequences, which are then validated for affinity [7]. However, obtaining high-performance aptamers from the SELEX process remains challenging due to insufficient library diversity, PCR bias, and low library enrichment efficiency [8–11]. The most frequently enriched sequences often lack the highest affinity and specificity [12]. Therefore, in silicon methods are needed to facilitate the discovery of high-performance aptamers.

Bioinformatics methods based on molecular docking are used in aptamer development, focusing on structural simulations [13, 14]. A typical workflow includes four main steps: secondary structure prediction, tertiary structure optimization, molecular docking with the target molecule, and molecular dynamics simulation of the docked complex [15, 16]. Secondary structure prediction tools include RNAfold [17], Mfold [18], and Vfold2D [19]. Tertiary structure prediction utilizes RNAComposer [20], 3dRNA [21], and Vfold3d [22]. Molecular docking simulations employ ZDOCK [23], AutoDock [24], AutoDock Vina [25], and NPDock [26], while AMBER [27] and GROMACS [28] are used for molecular dynamics simulations. Current docking methods cannot be used in analyzing large HT-SELEX data and struggle with predicting tertiary structures for long aptamers [16].

In recent years, the rapid development of machine learning has greatly enhanced aptamer development. Machine learning methods for aptamer development are broadly divided into two categories: clustering evaluation based on primary sequences and on secondary structures. Primary sequence clustering treats sequences as simple strings composed of nucleotides, performing clustering analysis as seen in AptaCluster [29], FASTAptamer [30], and FASTAptamer2 [31]. However, these models lack structural information, reducing accuracy since some aptamers are structurally conserved despite significant sequence variations, vice-versa. Secondary structure-based clustering focuses on the structural characteristics of local regions. For example, AptaTrace [32], Fast String-Based Clustering (FSBC) [33], RaptRanker [34] etc.

Deep learning analyzes relationships within data through deep neural networks that mimic human neurons. Di Gioacchino et al. created a Restricted Boltzmann Machine (RBM) model using thrombin HT-SELEX data demonstrating that log-likelihood scores are positively correlated with sequence fitness [35]. Iwano et al. designed a deep neural network RaptGen, based on a Variational AutoEncoder (VAE), which uses a Convolutional Neural Network (CNN) as an encoder and a Hidden Markov Model (Profile-HMM) as a decoder to generate and predict potential candidate aptamers [36]. Deep learning can enhance aptamer binding predictions but is limited by scarce training data, lack of labeled experiments, and costly procedures. The interpretability of these networks also needs improvement. During SELEX, sequence and substructural changes occur, influencing the population’s fitness landscape [37, 38].

In this study, we used rapamycin-specific aptamers isolated by our laboratory and thrombin-specific aptamers in a published work to evaluate the performance of our new developed APV-Sankey Toolbox. Rapamycin aptamers were obtained through HT-SELEX Data from <u>Rapa</u>mycin-Round X (RAPA-RX), where X stands for the number of rounds. By utilizing this advanced tool for systematic aptamer analysis and visualization, we successfully identified aptamers with strong binding affinity to rapamycin, demonstrating the efficacy of the APV-Sankey Toolbox in optimizing aptamer selection (see result section). Furthermore, the APV-Sankey Toolbox has been applied to the analysis of thrombin-specific aptamers (the results of aptamers of thrombin are mostly displayed in the supplement materials) that have been previously reported [39] [35]. Through the analysis of disparate datasets, the APV-Sankey Toolbox has demonstrated its ability to elucidate the differences in K-mer variations across rounds of specific SELEX methods.

### Implementation

APV-Sankey toolbox uses Python command functions, offering a comprehensive set of features to help users understand the evolutionary and structural characteristics of aptamers. **Figure 1** illustrates all the functions provided by the APV-Sankey toolkit. It allows for full sequence analyses to track evolutionary patterns in individuals or populations and monitors nucleotide ratios to evaluate interactions with targets. The toolbox enables users to identify the most common sequences across selection rounds using bar and line plots, while also allowing to discover potential functional small K-mers to discover recurring motifs and their binding properties. By evaluating K-mer frequencies, users can assess the abundance of common motifs. Hierarchical clustering methods can reveal relationships within aptamer populations, and dendrograms generated from Levenshtein edit distances can help to measure sequence differences. Additionally, users can concatenate overlapping K-mer segments to identify complete binding motifs and simulate K-mer evolution to explore evolutionary paths. Most importantly, users can create Sankey charts to illustrate relationships between stages of aptamer analysis, providing clarity in the screening process.

**Figure 1.**
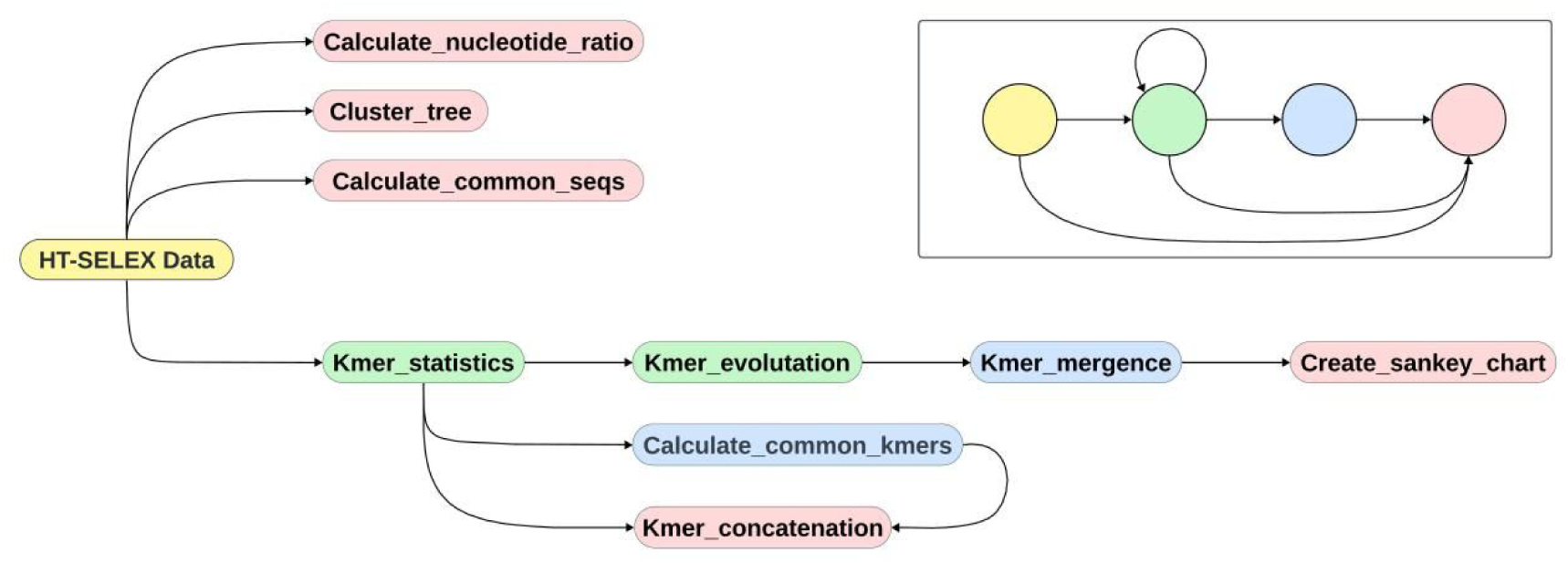
Diagram of functional connections within the APV-Sankey Toolbox. All pipelines must originate from HT-SELEX Data (yellow). Green modules represent intermediate steps within the pipeline and can feed into green modules, blue modules, or pink modules. Blue modules also represent intermediate steps but can only feed into pink modules, which always serve as the terminal steps of the pipeline. All black lines in the diagram are unidirectional. The simplified legend in the upper right corner indicates the directionality of the connections.

All methods and codes within the APV-Sankey Toolbox are available for free download on Gitee at the following repository: https://gitee.com/Zyyy202/APV-Sankey. Additional details can be found in the Supplementary Materials.

## Results

### Sequence-Level Individual/Population Tracking

Using HT-SELEX sequencing data of screening aptamers for Rapamycin as an example, we analyzed full sequences to track how individuals and populations evolved. The ***Calculate_common_seqs*** function outputs a figure with a bar plot displaying the counts of the top 10 most frequent common sequences in RAPA-R12 and RAPA-R15, and a line plot showing the count ranks of these sequences across the respective rounds (**Figure 2A**). This help to find out the most prevalent common sequences, likely of significant sequence-level importance. Monitoring these ratios across different selection rounds can reveal shifts in nucleotide composition indicating selective pressures or evolutionary trends within the aptamer population. Results of screening aptamers for thrombin and more explanations can be found in supplement materials.

**Figure 2.**
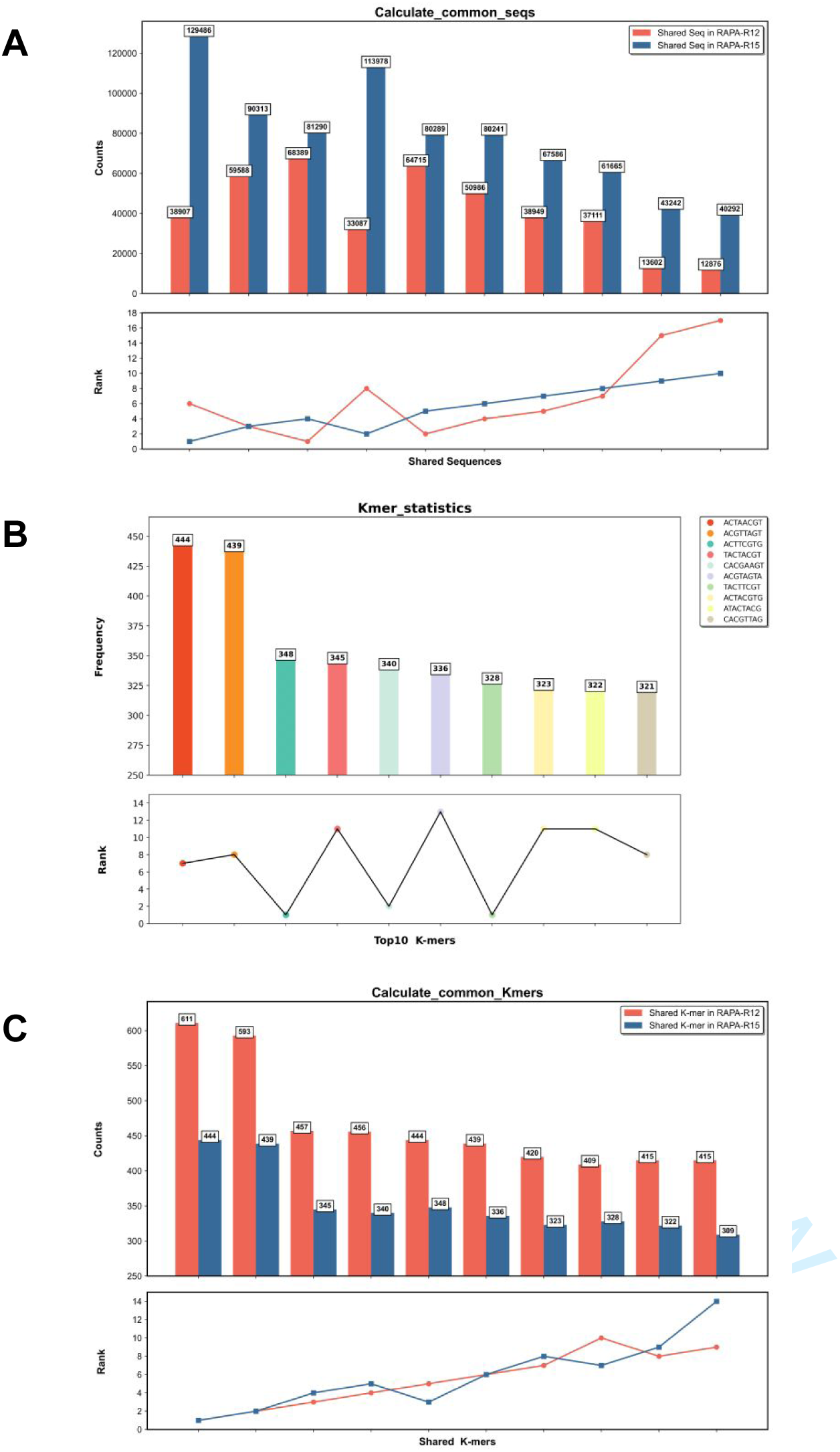
(A): The top 10 most frequent common sequences using RAPA-R12 and RAPA-R15 using Calculate_common_seqs method. The bar chart above illustrates the frequency of these common sequences in respective rounds, while the line chart below emphasizes their ranking in count for the corresponding round (the same to the following figures). (B): The top 10 most frequent K-mers using RAPA-R15 with the Kmer_statistics method (sequence counts are not considered). (C): The top 10 most frequent common K-mers using RAPA-R12 and RAPA-R15 with the Calculate_common_Kmers method (sequence counts are not considered).

Tracking the evolutionary patterns of individual sequences involves analyzing the sequences from a single selection round. This approach identifies high frequency sequences and assesses sequence diversity within that round. In contrast, population tracking spans of multiple selection rounds, providing a broader view of how the fitness landscapes evolve over time.

### Substructural Motif-Level Individual/Population

We applied the ***Kmer_statistics*** function to RAPA-R15 with K=8. The results showed the top 10 K-mers along with the highest-count sequences containing these K-mers (**Figure 2B**). Subsequent fluorescence competitive binding experiments confirmed that 3 out of the 6 sequences have affinity for rapamycin (**Figure S1**). Analyzing substructural K-mers within sequences to identify and track the evolutionary patterns of individuals or populations uncovers recurring motifs, provides insights into binding affinity, specificity, and overall structural properties, and enhances understanding of the evolutionary dynamics and functional characteristics of aptamer sequences.

Based on the single-round results from the ***Kmer_statistics*** method, we use the ***Calculate_common***_***Kmers*** method to identify shared K-mers across multiple rounds. This approach processes a list of K-mer statistics files and outputs a .xlsx file summarizing the common K-mers, their frequencies, and ranks across the input files, offering a comprehensive view of conserved motifs. **Figure 2C** presents the frequency and ranking of the top 10 most abundant shared K-mers in RAPA-R12 and RAPA-R15 across the corresponding rounds. Results are visualized as bar charts and line graphs to depict the enrichment of K-mers throughout the SELEX process. This analysis pinpoints consistently appearing K-mers across different rounds, which may highlight key functional elements. Additionally, the dominant K-mers identified are further visualized in Sankey charts (see ***Create_sankey_chart*** section), providing insights for potential truncation or chemical modification in nucleic acid engineering.

### Hierarchical Clustering of Aptamer Sequences

The ***Cluster_tree*** method supports optional affinity values, generating dendrogram bar plots with showing affinity information. This dual analysis offers insights into sequence similarities and their affinities. **Figure 3** shows the hierarchical clustering of candidate sequences from RAPA-R15, where affinity was evaluated via fluorescence competitive binding experiments. Notably, high-affinity sequences R15-8, R15-56, and R15-2 clustered together, highlighting functional regions within the clusters. This method provides a robust framework for classifying and understanding aptamer sequences, deepening insights into their evolutionary and functional dynamics.

**Figure 3.**
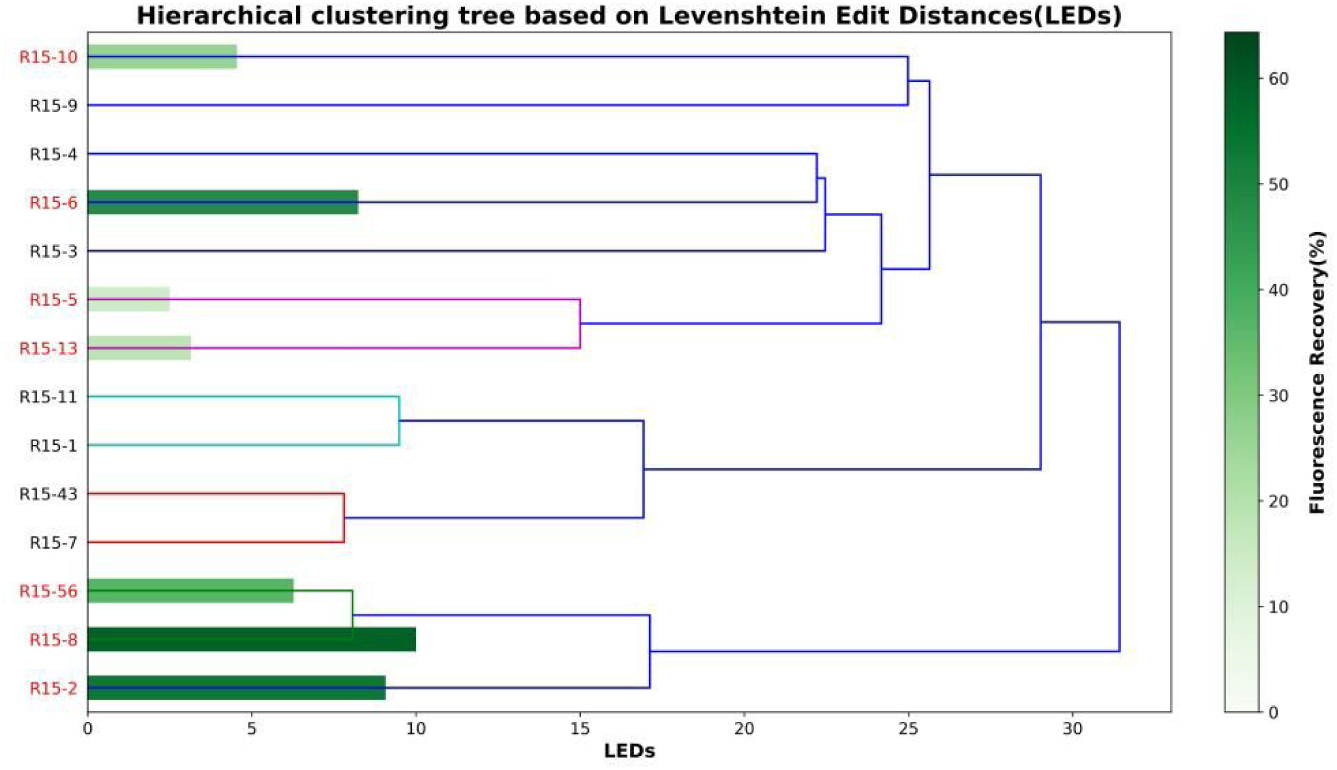
Hierarchical clustering tree generated by the Cluster_tree method to a subset of candidate sequences from RAPA-R15. Red labels indicate sequences with measured affinity, while the green bar plots represents the relative magnitude of fluorescence recovery values.

### K-mer Concatenation

Traditional K-mer analysis often results in fragmented information, making it challenging to derive meaningful insights. The ***Kmer_concatenation*** method starts with single-round multi-round common K-mers and iteratively concatenates K-mer pairs until no further concatenation is possible or only one longest segment remains (**Figure S4**). By concatenating overlapping K-mers, this method consolidates information to identify complete binding motifs. **Table 1** shows the application of the ***Kmer_concatenation*** method to RAPA-R12, RAPA-R15, and their common K-mers. Despite different initial K-mer lengths, the longest concatenated segments (LCS) consistently converge into a few distinct sequences, predicting high-affinity aptamers and demonstrating library enrichment. The consistency of results across different rounds affirms the robustness of the method.

**Table 1.**
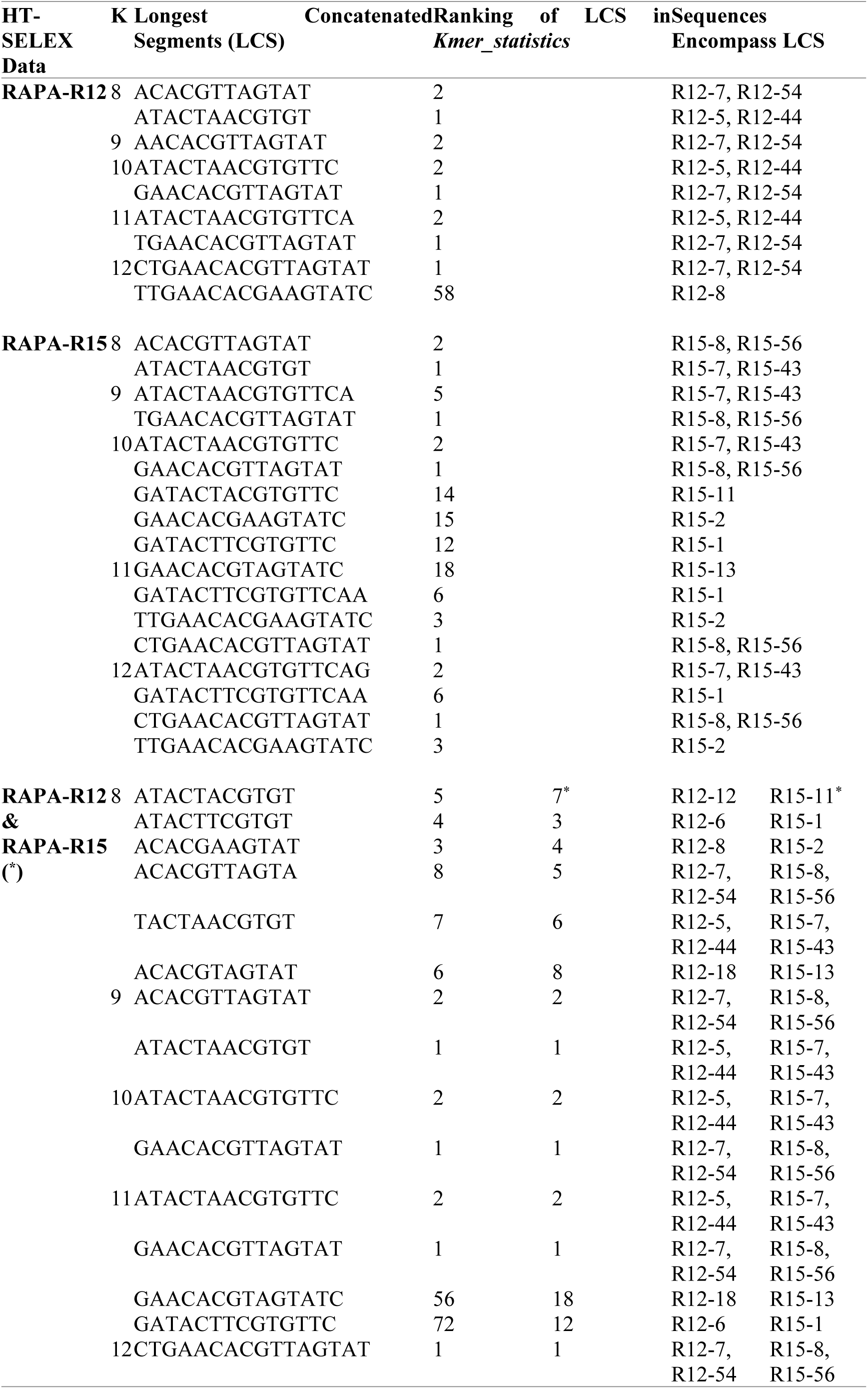
Results of Applying *Kmer_concatenation* method on K-mers in RAPA-R12, RAPA-R15, and their shared K-mers.

### K-mer Evolution & K-mer Mergence

The ***Kmer_mergence*** method processes result from the ***Kmer_evolution*** method by merging duplicates and removing K-mers with decreasing frequencies to eliminate redundancy. This simplifies evolutionary patterns, enhances data clarity, and supports structured Sankey chart creation.

### Create Sankey chart

Aptamers fold into specific tertiary structures to bind target molecules, with only certain nucleotides significantly impacting binding [40]. While some K-mer base changes have minimal structural effect, others can lead to functional loss or reduced target binding. K-mers may become dominant in later rounds, be eliminated early, or persist throughout the process. Visualizing their progression is crucial for understanding the dynamic evolution of fitness landscapes. The ***Create_sankey_chart*** method generates intuitive Sankey charts to visualize relationships between rounds of aptamer analysis, enhancing understanding of the SELEX screening process. This method enables researchers to track K-mer trajectories, pinpoint key transition points, and make data-driven decisions to refine the SELEX process for optimal aptamer selection.

**Figure 4** illustrates a five-layer Sankey chart from RAPA-R5, RAPA-R7, RAPA-R8, RAPA-R12, and RAPA-R15 data. Each column represents a SELEX round, with the height of rectangular boxes indicating K-mer frequencies, and colors representing the corresponding K-mers. Flow lines between columns display evolutionary transitions. ***Create_sankey_chart*** provides a comprehensive view of K-mer selection and enrichment, highlighting conserved sites and evolutionary trends [41, 42]. Using the ***Kmer_evolution*** and ***Kmer_mergence*** methods for preprocessing, it generates interactive Sankey charts. Each node represents a K-mer, with unique colors and flow lines showing evolutionary transitions and quantity changes. Researchers can explore the data through simple hover and click actions, offering a clear view of K-mer dynamics and structure.

**Figure 4.**
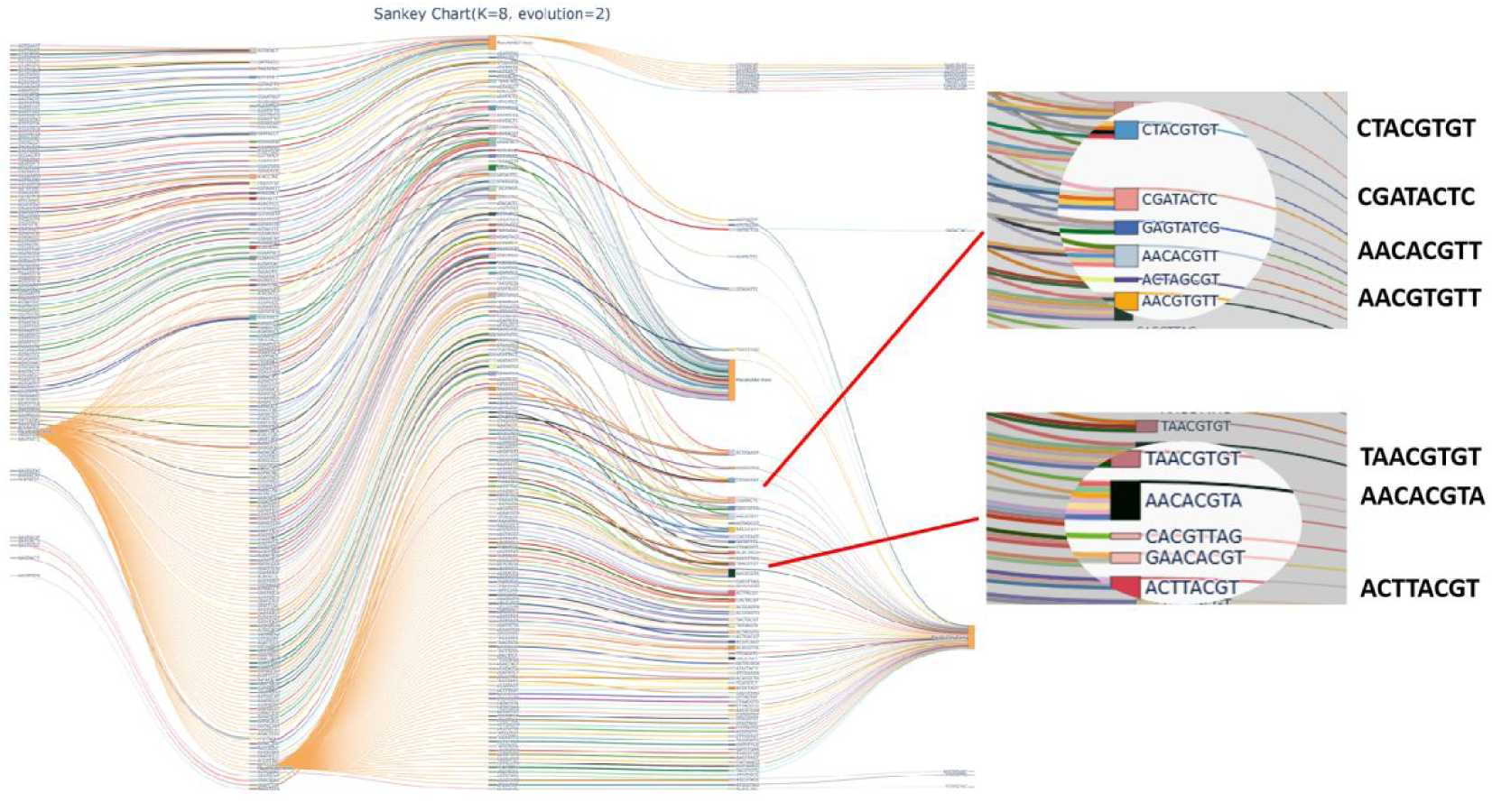
A five-layer Sankey chart constructed using RAPA-R5, RAPA-R7, RAPA-R8, RAPA-R12, and RAPA-R15.

## Discussion

APV-Sankey’s key innovation is the use of Sankey charts for visualizing the flow and relationships of sequences across SELEX rounds, offering an intuitive representation of transitions and relationships. This innovation addresses the limitations of theoretical research in SELEX, which has often relied on empirical parameter settings. SELEX technology has evolved over the past 30 years, theoretical research on SELEX itself has been limited, often relying on experience-based parameter settings [43]. Different SELEX methods are used depending on the target type, with recent interest shifting from protein-targeting methods like MB-SELEX [44] and CE-SELEX [45] to those for small molecule targets, such as homogenous-phase GO-SELEX and solid-phase Capture-SELEX [46, 47]. SELEX technology is diverse, with significant differences in wet lab parameters across techniques, often lacking theoretical guidance. Sankey charts provide quantitative insights into sequence abundance changes, aiding in the optimization of experimental parameters and the development of more effective screening methods.

Additionally, PCR amplification errors, which mimic biological evolution’s mutation process, vary in frequency. Unlike random library screening, mutations in later rounds occur in retained sequences, potentially generating higher-affinity aptamers. Sankey diagrams can capture these evolutionary patterns, offering a unique perspective on understanding and harnessing these variations in the selection process. APV-Sankey effectively traces and reveals the evolutionary trends of individual and population sequences which is crucial for optimizing candidate sequences through screening truncation optimization for downstream applications [48, 49].

Unfortunately, most reported truncation methods simply describe the truncation approach without providing specific evidence, which may lead to some loss of sequences functionality. Traditional methods may affect the accuracy of nucleic acid engineering due to the absence of detailed tracing. APV-Sankey addresses this by enabling users to track the entire SELEX process, offering dynamic interactive features that simplify tracing tasks for both novices and experts. Visualizing results with Sankey charts also provide evidence for aptamer truncation in subsequent nucleic acid engineering.

APV-Sankey provides comprehensive insights into sequence site mutations during tracing. It enables tracking of individual sequence sites changes. Directed changes and chemical modifications of specific sites can significantly enhance enzyme degradation resistance and improve the affinity and specificity of aptamers [50]. By providing detailed views of sequence site changes, APV-Sankey helps researchers make informed decisions on directed modifications, improving enzyme resistance, affinity, and specificity of aptamers to provide novel insights for the high-performance screening of nucleic acid aptamers.

## Conclusions

The APV-Sankey Toolbox is a specialized toolbox designed to enhance the efficiency of aptamer research through advanced sequence analysis and visualization. It supports key tasks including sequence data analysis, similarity calculations, plotting, and clustering. The toolbox allows for the optimization of algorithms tailored to specific aptamer sequences, ensuring both efficient and accurate analysis.

The Toolbox uses Sankey diagrams providing a clear and intuitive method for visualizing relationships between different stages of the SELEX process. This enhances the interpretability of results, enabling researchers to explore sequencing libraries more deeply and identify high-affinity sequences. The APV-Sankey Toolbox also offers objective data on K-mer mutation tracing, SELEX method selection, and aptamer trimming and modification. These features make it a valuable resource for advancing research in molecular biology, diagnostics, and therapeutics related to aptamers.

In summary, the APV-Sankey toolbox not only accelerates the pace of aptamer-related research but also contributes to the broader scientific community by providing a robust and innovative toolkit for the exploration of aptamer sequences.

## Key Points

- We developed APV-Sankey, a novel and versatile toolbox designed for the rapid aptamer screening and visualization of the enrichment process.
- APV-Sankey uses Sankey charts for interactive and informative visualizations, facilitating tracing the evolution of K-mers across SELEX rounds, aiding in the identification and selection of high-affinity sequences containing K-mers while excluding high-frequency sequences without affinity.
- We successfully identified high-performance candidate aptamers of both rapamycin and thrombin using APV-Sankey.

## Acknowledgement

This work was supported by National Natural Science Foundation of China (22374104, 21705112), Capacity Building for Sci-Tech Innovation-Fundamental Scientific Research Funds (KM201910028014), Beijing outstanding talents training fund for youth backbone individual project (2017000020124G081).

## Supplementary Information

## 1. Getting Started

This guide is organized into three sections. **Section 2** provides specific information about the wet-lab experimental data used in our analyses, while **Section 3** offers an in-depth exploration of each functional module, complete with detailed step-by-step instructions and examples to guide users in effectively utilizing the toolbox. It is important to note that the current version of APV-Sankey Toolbox operates exclusively through command-line interfaces. To ensure ease of use, we provide command scripts necessary for running each function. We anticipate releasing a Graphical User Interface (GUI) version of the APV-Sankey Toolbox in the next update.

**Table S1** summarizes the input and output file types associated with each function in APV-Sankey. Users can choose to execute individual functions based on their specific needs or combine multiple functions to perform a comprehensive analysis workflow. It is important to note that APV-Sankey is focused on processing sequence data obtained from high-throughput sequencing (HT-SELEX data) and performing downstream analyses; therefore, it does not include any steps or methods for data preprocessing. When applying the APV-Sankey Toolbox to custom input data, the data format must meet the required specifications, as detailed in **Section 3.1**.

**Table S1.**
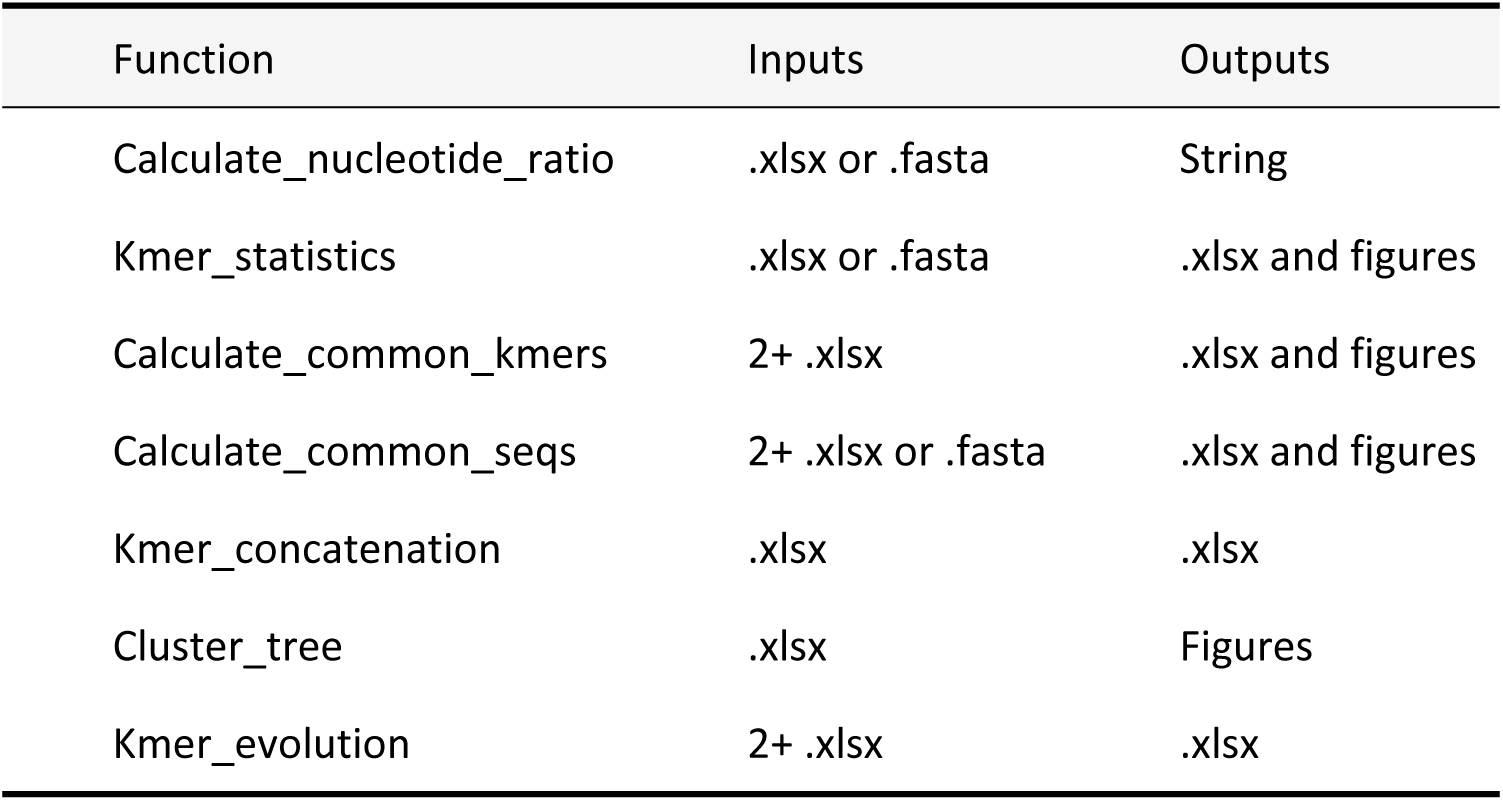
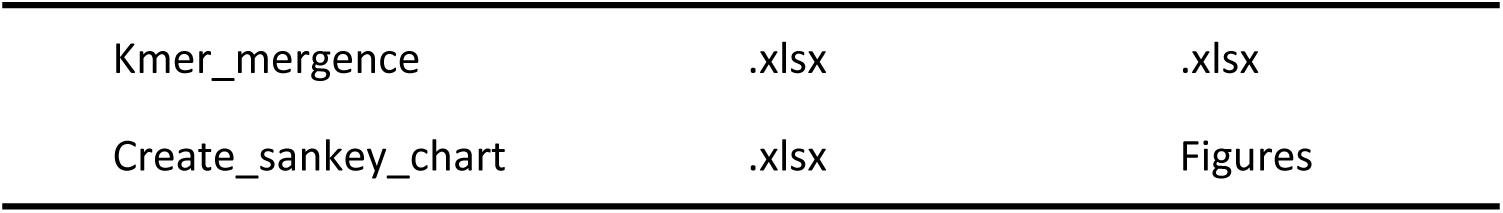
File Types for Each Functional Module in APV-Sankey.

| Function | Inputs | Outputs |
| --- | --- | --- |
| Calculate_nucleotide_ratio | .xlsx or .fasta | String |
| Kmer_statistics | .xlsx or .fasta | .xlsx and figures |
| Calculate_common_kmers | 2+ .xlsx | .xlsx and figures |
| Calculate_common_seqs | 2+ .xlsx or .fasta | .xlsx and figures |
| Kmer_concatenation | .xlsx | .xlsx |
| Cluster_tree | .xlsx | Figures |
| Kmer_evolution | 2+ .xlsx | .xlsx |
| Kmer_mergence | .xlsx | .xlsx |
| Create_sankey_chart | .xlsx | Figures |

## 2 Data

All data presented in this guide are divided into two main parts. The first part pertains to the data on <u>rapa</u>mycin-specific aptamers (RAPA), which were obtained through laboratory screening. In brief, we successfully isolated rapamycin-specific aptamers using the Capture-SELEX method with agarose beads, completing 19 rounds of selection. **Section 2.1** provides a detailed explanation of how the Capture-SELEX process was conducted in our laboratory, while **Section 2.2** summarizes the sequence information used in the wet-lab experiments. The second set of data concerns the thrombin-specific aptamer library, with further details could be seen in (Zhou, et al., 2019) and (Di Gioacchino, et al., 2022).

### 2.1 Experimental Procedure

#### 2.1.1 Capture-SELEX

1. Rounds 1-14: No negative selection was applied. The library and capture strand were mixed in selection buffer, heated, and cooled to allow hybridization. Streptavidin-coated agarose beads were washed, and the hybridized complex was added. After multiple washes to remove unbound sequences, the library was eluted for positive selection.
2. Rounds 14-19: Same as rounds 1-14, but with an additional negative selection using cyclosporine after the washes. The negative selection eluate was collected.
3. ssDNA Preparation: Positive selection eluates underwent PCR, and the product was processed with streptavidin-coated agarose beads to isolate ssDNA. The ssDNA was eluted, quantified, and stored for further use.

In total, 19 selection rounds were performed, and PCR products from selected rounds were prepared for high-throughput sequencing by GENEWIZ.

#### 2.1.2 Fluorescence Competitive Binding Experiments of Candidate Aptamers

Affinity measurements for candidate aptamers were conducted using a fluorescence competition assay with full-length aptamers. The procedure involved dissolving 6 pmol of the aptamer and 30 pmol of the capture strand in selection buffer, heating, and cooling for hybridization. Rapamycin was added to a final concentration of 1 μM, and the mixture was incubated for 30 minutes. Fluorescence emission at 520 nm was measured using a spectrometer, and fluorescence recovery was calculated:

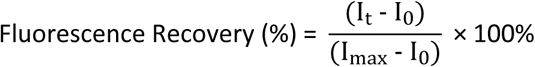

Where:

- I*_max_* is the fluorescence signal of 50 nM candidate aptamer
- I*_t_* is the fluorescence signal after incubating rapamycin in the candidate aptamer-capture strand hybridization system
- I*_0_* is the fluorescence signal of the candidate aptamer-capture strand hybridization system

This calculation allowed for the assessment of the binding affinity of the aptamer to rapamycin by observing changes in fluorescence emission.

### 2.2 Results Summary

**Table S2** provides information on the libraries, primers, and modified sequences used for the isolation of rapamycin-binding aptamers. **Figure S1** illustrates the results of the affinity assessment of R15 candidate aptamers using a fluorescence competitive assay.

**Table S2.**
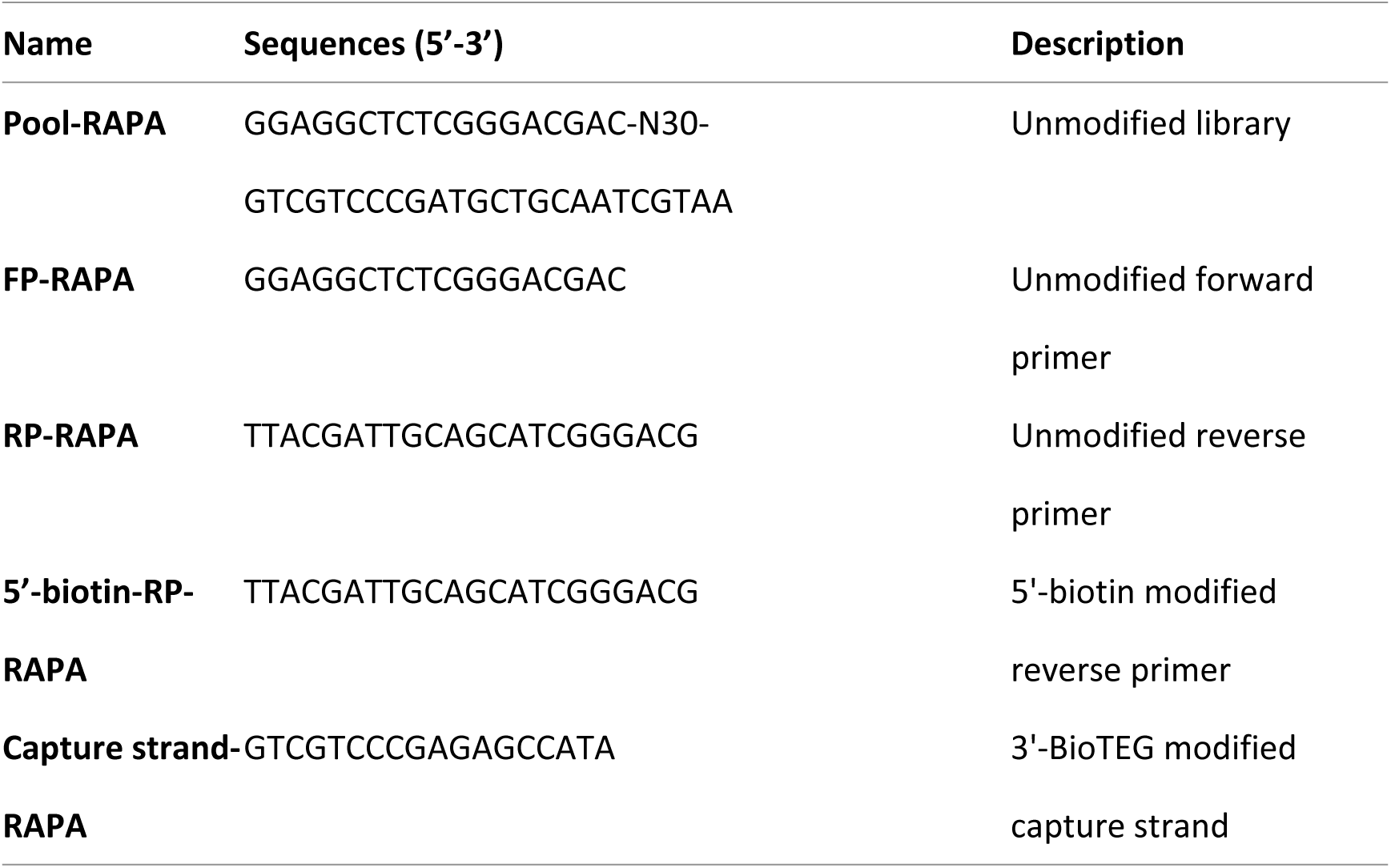
Initial Libraries, Primers, and Modified Sequences.

**Figure S1.**
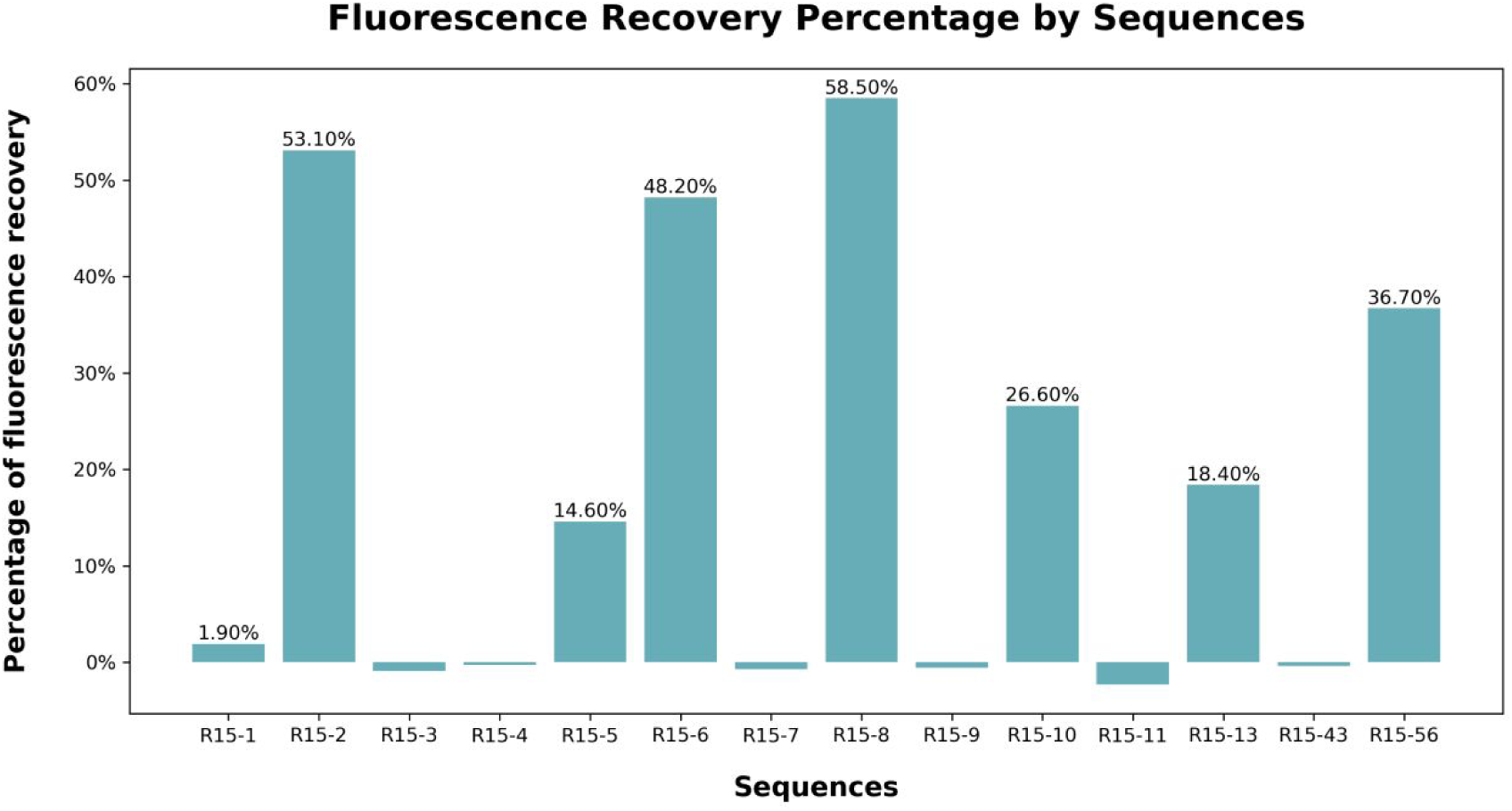
Results of the Fluorescence Competitive Assay of RAPA-R15 Candidate Aptamers

## 3 Methods

This Python code utilizes popular libraries such as NumPy, Pandas, Biopython, and Plotly to perform various bioinformatics tasks. While a graphical user interface (GUI) version of the APV-Sankey Toolbox is in development shortly, the current command-line version ensures broad accessibility, making it a valuable for both novice and experienced researchers in the aptamer community. All code has been tested on both workstations and personal laptops with the following specifications, so that users can easily apply APV-Sankey in any situation:

Workstation: CPU: Intel(R) Xeon(R) Gold 6128 CPU @ 3.40GHz; Memory: 64G; OS: Windows 10

Personal laptop: CPU: Intel(R) Core (TM) i7-14650HX CPU @ 2.20GHz; Memory: 8G; OS: Windows 11.

### Sankey diagram

In a commonly used Sankey diagram, the calculation of flows is typically based on the principle that the total amount of flow into and out of a node must be equal. However, in this work we used normalized data in each round in SELEX, so it means that in each round of flow transfer, the outflow from each node is normalized relative to the total inflow into that node. This way, the sum of the outflows from each node will equal 1 (or 100%).

#### 1. Calculate the Total inflow for Each Node

For each node i, first calculate the total inflow *F*_*in*,*i*_:

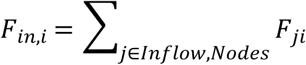

#### 2. Normalize the Outflow for Each Node

For each node i, normalize its outflow *F*_*ik*_:, where k is the outflow node:

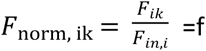

#### 3. Ensure the Normalized Outflow Sum Equals 1

For each node i, the sum of the normalized outflows should equal 1:

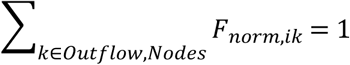

#### 4. Calculate the Normalized Flow Values

Convert the normalized flows back to actual flow values using the total inflow:

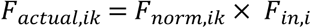

#### 5. Update the Flow in the Sankey Diagram

In the Sankey diagram, use Fctual,ik to represent the flow from node i to node k, so that the sum of the outflows from each node will equal the inflow.

#### 6. Repeat the Normalization Process

The Sankey diagram includes multiple rounds of flow transfer, so we repeat the normalization process for each round.

Normalization allows for a fair comparison of flows by adjusting them to a common scale (0 to 1 in this case). In the Sankey diagram, the width of the arrows is typically proportional to the flow values. With normalized flows, the width of the arrows will reflect the relative importance of each flow, rather than their absolute values. This approach provides flexibility in visualizing flows where the total inflow and outflow for each node may not be equal. It’s particularly useful in scenarios where the focus is on the distribution of flows rather than their absolute values.

### Data description

All data utilized in this article were obtained from previous laboratory studies. In simple terms, they are aptamer datasets targeting rapamycin, obtained through 19 rounds of selection using the Capture-SELEX method. High-throughput sequencing was performed on Round5 (RAPA-R5), Round7 (RAPA-R7), Round8 (RAPA-R8), Round12 (RAPA-R12), Round15 (RAPA-R15), Round17 (RAPA-R17), and Round19 (RAPA-R19). See Supplementary Materials for details.

We introduce the functions of the APV-Sankey Toolbox, providing a comprehensive operational guide to help users efficiently utilize the toolbox. This guide covers input data formats, command-line scripts, parameter details, method execution logic, and result interpretation. While we use datasets from our laboratory as examples, any data that meets the specified format and requirements can be used, and users can substitute their own data according to the instructions in **Section 3.1**.

### 3.1 HT-SELEX Data

The APV-Sankey Toolbox accepts preprocessed .fasta or .xlsx files as input. Users must provide high-throughput sequencing data in one of these formats, with .xlsx files including the columns in **Figure S2** and .fasta files following the format in **Figure S3**.

**Figure S2.**
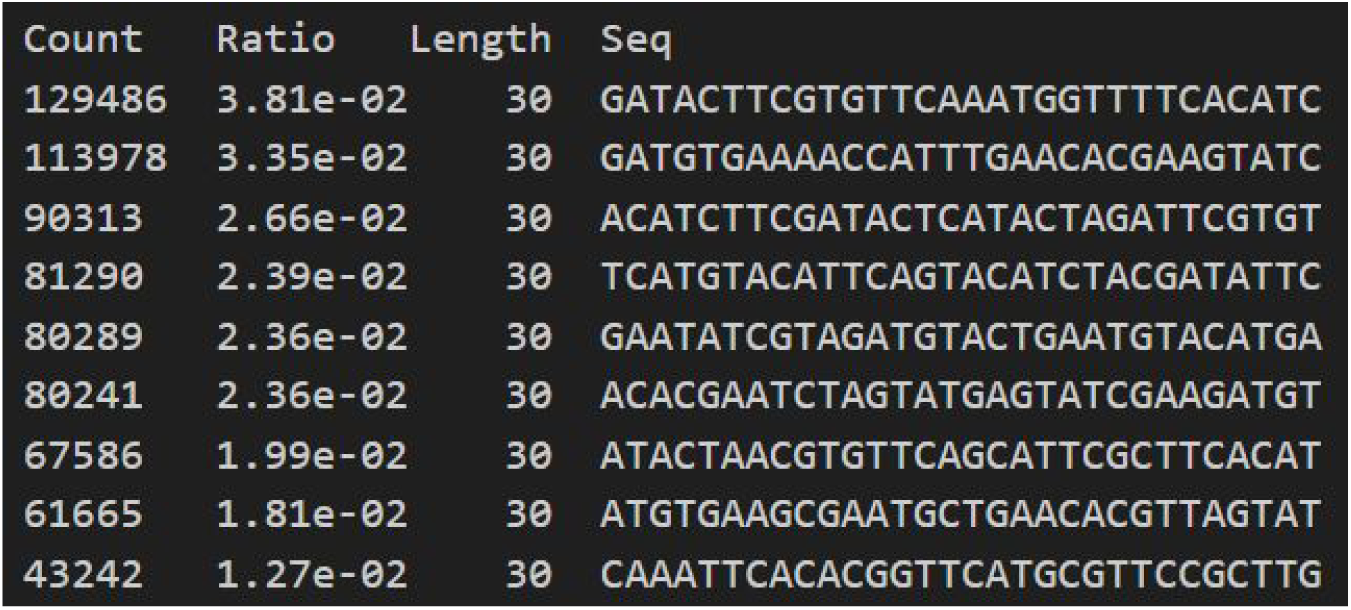
Input format for .xlsx files: ‘Count’ shows sequence count, ‘Length’ indicates sequence length, ‘Seq’ contains sequence data, and ‘Ratio’ represents sequence abundance (current counts/total library counts).

**Figure S3.**
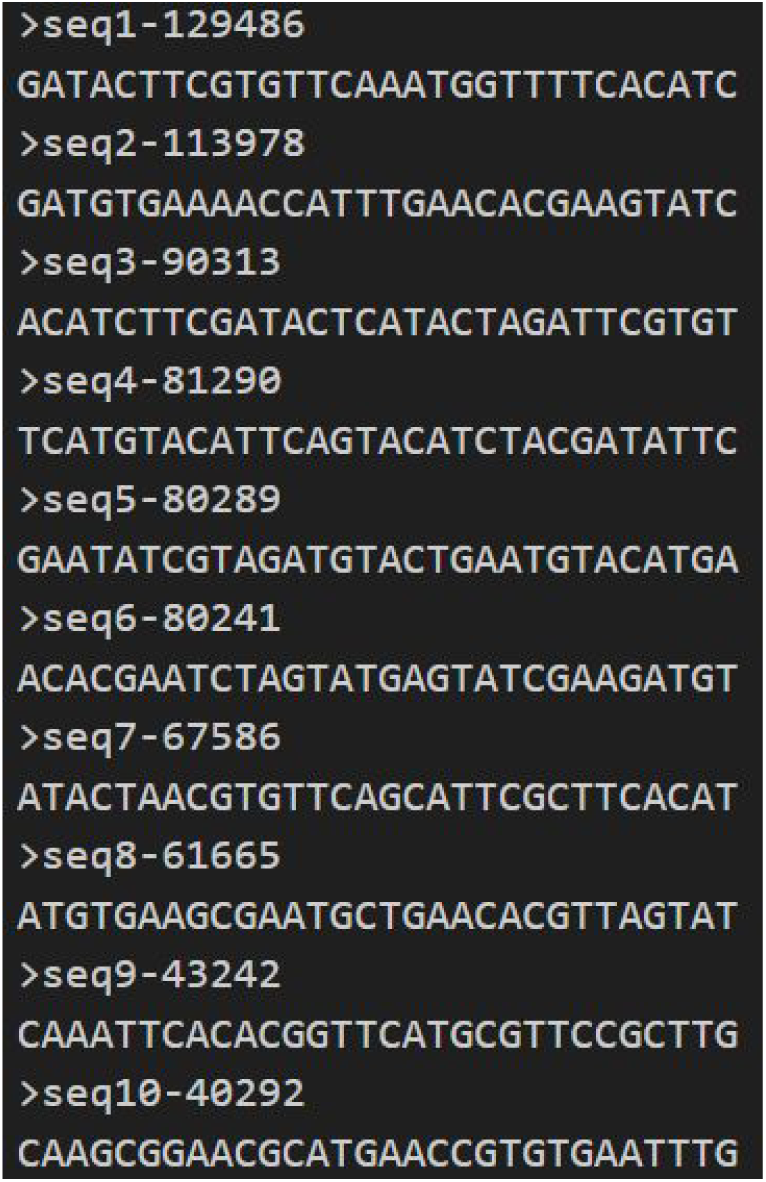
Input format for .fasta files: The first line starts with ‘>’, where ‘>seqX-Y’ indicates ID (X) and counts (Y). The second line contains the sequence.

### 3.2 Calculate_nucleotide_ratio

The ***Calculate_nucleotide_ratio*** quantifies nucleotide composition in .xlsx or .fasta sequence files. It computes the ratios of adenine (A), guanine(G), cytosine (C), and uracil(U) (for RNA) or thymine (T)(for DNA), based on the ‘is_rna’ parameter.

A detailed description of the method’s application is as follows:

- Calculate_nucleotide_ratio >>> Description - Calculate the nucleotide ratio of the given sequences and print the result >>> Parameters - filepath : string (‘/path/to/*.xlsx or .fasta’) is_rna (default: False) : bool >>> Returns – None >>> Examples - Calculate_nucleotide_ratio(‘/path/to/sequences.fasta’, is_rna=True)

### 3.3 Kmer_statistics

The ***Kmer_statistics*** method analyzes K-mer frequency distribution in nucleotide sequences, helping identify recurring patterns and their abundances. It processes .xlsx or .fasta files and allows users to specify the K-mer length (‘K’) and the number of top K-mers (‘n_Kmers’) for output. An optional parameter, ‘weigh_sequences_by_count,’ weights K-mer frequencies based on sequence counts from .xlsx files or sequence headers in .fasta files.

The ***Kmer_statistics*** reads the input file, extracts sequences and counts, and calculates K-mer frequencies. It sorts K-mers by frequency to identify the top n_Kmers and records their occurrences, sequences, and counts. The output includes K-mer, frequency, and associated sequences and counts.

A detailed description of the method’s application is as follows:

- Kmer_statistics >>> Description - Perform K-mer statistics and output the results to an Excel file >>> Parameters - filepath : string (‘/path/to/*.xlsx or .fasta’) K : int (The length of the K-mers to analyze) n_Kmers : int (The number of top K-mers to include in the output) output_filepath : string (‘/path/to/*.xlsx) weigh_sequences_by_count (default: False) : bool >>> Returns - An Excel file A plot image >>> Examples - Kmer_statistics(‘/path/to/sequences.fasta’, 3, 5, ‘/path/to/output.xlsx’, weigh_sequences_by_count=True)

### 3.4 Calculate_common_kmers

The ***Calculate_common_kmers*** method identifies conserved sequence motifs by analyzing K-mer frequencies across multiple datasets. It processes K-mer statistics Excel files and outputs an Excel file with common K-mers, their frequencies, and ranks across all input files.

The method loads K-mer data from each file, finds common K-mers, and calculates their frequency and rank in each dataset. It also computes the total frequency of each common K-mer across all datasets. The output file includes the K-mer, its frequency, rank, and total count across all files.

A detailed description of the method’s application is as follows:

- Calculate_common_kmers >>> Description - Calculate the common K-mers and their frequencies across multiple files >>> Parameters - common_kmers_input : list of string ([‘/path/to/file1.xlsx’, ‘/path/to/file2.fasta’]) common_kmers_output : string (/path/to/*.xlsx) >>> Returns - An Excel file A plot image >>> Examples - Calculate_common_kmers([‘/path/to/file1.xlsx’, ‘/path/to/file2.xlsx’], ‘/path/to/output.xlsx’)

### 3.5 Calculate_common_seqs

The ***Calculate_common_seqs*** method identifies and quantifies common subsequences across multiple aptamer datasets. It processes sequence statistics from Excel (.xlsx) or FASTA (.fasta) files and outputs an Excel file with the common sequences and their frequencies.

The method extracts sequences and counts from .xlsx or parses them from .fasta files. It compiles individual and total counts for each common sequence across all files. The output includes the sequence, counts from each file, and the total count, sorted by frequency.

A detailed description of the method’s application is as follows:

- Calculate_common_seqs >>> Description - Calculate the common sequences and their counts across multiple files >>> Parameters - common_seqs_input : list of string ([‘/path/to/file1.xlsx’, ‘/path/to/file2.fasta’]) common_seqs_output : string (‘/path/to/*.xlsx’) >>> Returns - An Excel file A plot image >>> Examples - Calculate_common_seqs([‘/path/to/file1.xlsx’, ‘/path/to/file2.xlsx’], ‘/path/to/output.xlsx’)

### 3.6 Kmer_concatenation

The ***Kmer_concatenation*** method assembles K-mers based on overlapping regions to form longer sequences, helping identify binding regions. For instance, if we have the K-mers ‘GCATCCTG’, ‘ATCCTGAC’, ‘CCTGACGG’, and ‘ACGGTACC’, which can be viewed as fragments of a longer sequence ‘GCATCCTGACGGTACC’, the method will concatenate these K-mer fragments to reconstruct the longer sequence. This is particularly useful for identifying the longest complete potential binding units.

The method reads K-mers from an input Excel file, combines overlapping K-mer pairs, and repeats until no further concatenation is possible. It saves the final concatenated sequences and the K-mers used in each step to an output Excel file.

A detailed description of the method’s application is as follows:

- Kmer_concatenation >>> Description - Concatenate K-mers based on certain rules and save the results to an Excel file >>> Parameters - kmer_concatenation_input : string (‘/path/to/*.xlsx’) kmer_concatenation_output : string (‘/path/to/*.xlsx’) sheet_name : string (The name of the sheet in the input Excel file) >>> Returns - An Excel file >>> Examples - Kmer_concatenation(‘/path/to/input.xlsx’, ‘/path/to/output.xlsx’, ‘Sheet_name=0’)

### 3.7 Cluster_tree

The ***Cluster_tree*** generates hierarchical clustering trees (dendrograms) based on sequence similarities using Levenshtein Edit Distances (LEDs). It optionally includes ‘Affinity’ values for additional insights.

The method reads sequences and IDs from an Excel file, calculates the LED matrix, and generates the dendrogram. If ‘Affinity’ values are included, they are visualized with a heatmap and bar plots alongside the dendrogram, showing both sequence similarities and affinities.

A detailed description of the method’s application is as follows:

- Cluster_tree >>> Description - Generate a clustering dendrogram based on Levenshtein Edit Distances (LEDs) between sequences and/or ‘Affinity’ values >>> Parameters - cluster_filepath : string (‘/path/to/*.xlsx’) The file may optionally contain an ‘Affinity’ column with percentage values >>> Returns - A clustering dendrogram plot with optional ‘Affinity’ annotations >>> Examples - Cluster_tree(‘/path/to/file.xlsx’)

### 3.8 Kmer_evolution

The ***Kmer_evolution*** method compares K-mers between consecutive files, identifying when a K-mer has evolved based on a specified maximum number of changes (parameter ‘evolution_count’). It reports raw and evolved K-mers, their frequencies, and mutation details.

The method loads K-mer data from each file, checks if K-mers in file1 differ from those in file2 by the allowed changes, and identifies them as evolved. It compiles the results, including frequencies and mutation details, into an Excel file, removing duplicates for clarity.

A detailed description of the method’s application is as follows:

- Kmer_evolution >>> Description - Perform evolution analysis on K-mers between multiple files >>> Parameters - kmer_evolution_input : list of string ([‘/path/to/file1.xlsx’, ‘/path/to/file2.xlsx’]) evolution_count (default = 1): The maximum number of evolutions allowed for a K-mer to be considered as evolved kmer_evolution_output : string (‘/path/to/*.xlsx’) >>> Returns - An Excel file >>> Examples - Kmer_evolution([‘/path/to/file1.xlsx’, ‘/path/to/file2.xlsx’], evolution_count=1, output_file=’/path/to/output.xlsx’)

### 3.9 Kmer_mergence

The ***Kmer_mergence*** method merges overlapping K-mers between adjacent groups from an input Excel file, ensuring each unique sequence is represented once.

It reads the input file, identifies overlapping K-mers, and combines them, aggregating their frequency data. The method outputs an Excel file with the merged K-mers, their total frequencies, and relevant group information. It also removes rows with null or zero values to ensure a clean and meaningful final dataset.

A detailed description of the method’s application is as follows:

- Kmer_mergence >>> Description - Merge K-mers between adjacent groups in an input Excel file and save the results to an output Excel file >>> Parameters - kmer_mergence_input : string (‘/path/to/*.xlsx’) kmer_mergence_output : string (‘/path/to/*.xlsx’) >>> Returns - An Excel file >>> Examples - Kmer_mergence(‘/path/to/input.xlsx’, ‘/path/to/output.xlsx’)

### 3.10 Create_sankey_chart

The ***Create_sankey_chart*** generates Sankey charts to visualize K-mer evolution and frequency distributions across SELEX rounds. It uses data from the ***Kmer_mergence*** method, linking nodes dynamically based on K-mer presence and frequency across consecutive rounds, making it ideal for tracking K-mer flow and evolution.

The method reads an input Excel file (parameter ‘sankey_input’) containing worksheets with K-mers, frequencies, and evolutionary data for each round. K-mers are extracted as nodes, with new or evolving sequences added. Nodes are uniquely stored and color-coded for clarity, and links between nodes are weighted by frequency to depict K-mer flow. Plotly is used to generate the chart, displaying node labels, colors, and link values, offering a clear visualization of K-mer evolution through SELEX rounds.

A detailed description of the method’s application is as follows:

- Create_sankey_chart >>> Description - Create a Sankey chart based on K-mers from an Excel file and display it >>> Parameters - sankey_input : string (‘/path/to/*.xlsx’) >>> Returns - A Sankey chart >>> Examples - Create_sankey_chart(‘/path/to/data.xlsx’)

## 4. Operating notes

1. HT-SELEX Data: The pipeline accepts preprocessed .fasta or .xlsx libraries as input.

High-Throughput Systematic Evolution of Ligands by Exponential Enrichment (HT-SELEX) is a powerful method for selecting high-affinity nucleic acid aptamers from large combinatorial libraries. The APV-Sankey analysis pipeline accepts single-round high-throughput sequencing libraries in .fasta or .xlsx formats as inputs. This serves as the starting point for all downstream processes, assuming that necessary data preprocessing has been completed beforehand. Preprocessing tasks include unique sequence frequency calculation, quality-based sequence filtering, and removal of sequencing adapters or linkers. Users are required to perform these preprocessing steps prior to using tools such as FastAptamer2.0, fastp and FASTX-Toolkit. This foundational step ensures that all downstream processes operate on well-defined and preprocessed ***HT-SELEX Data***.

The ***Kmer_statistics*** method analyzes K-mer frequency distributions within nucleotide sequences to identifiing recurring motifs and their abundance. By specifying the K-mer length (K) and the number of top K-mers to include in the output (n_Kmers), the method offers flexibility for detailed analysis. An optional parameter ‘weigh_sequences_by_count’ allows for weigh K-mer frequencies based on sequence counts, ensuring that the analysis accurately reflects the true abundance of each K-mer in the HT-SELEX data.

Traditional K-mer analysis often results in fragmented information, making it challenging to derive meaningful insights. By reconstructing a longer K-mer from several shorter K-mers, researchers can identify complete binding motifs that may not be apparent from individual K-mer alone. The ***Kmer_concatenation*** method systematically concatenates K-mers based on overlapping regions, simulating the formation of longer motifs from shorter fragments. For instance, given four K-mers ‘GCATCCTG’, ‘ATCCTGAC’, ‘CCTGACGG’, and ‘ACGGTACC’, each can be viewed as a fragment of a longer K-mer ‘GCATCCTGACGGTACC’. This method is particularly useful for identifying the longest complete potential binding units, offering a holistic view of the evolution of fitness landscapes within the sequences.

The ***Kmer_concatenation*** method starts with single-round multi-round common K-mers and iteratively concatenates K-mer pairs until no further concatenation is possible or only one longest segment remains. This automated process produces an output file detailing the concatenation results and the K-mers used at each step, allowing for detailed tracking.

### Kmer_evolution & Kmer_mergence

Simulates Tracing K-mer evolutions to explore evolutionary trajectories within fitness landscapes, simplifing and integrating the K-mer information.

Exploring the evolutionary dynamics of K-mers in aptamers is crucial for understanding how sequences enrich towards high affinity. By simulating the evolution of K-mer, researchers can trace evolutionary pathways and reveal trajectories within the fitness landscapes.

The ***Kmer_evolution*** methodidentifies evolved K-mers and quantify their changes, across multiple rounds, tracking their characteristics over successive SELEX stages This analysis reveals the selection pressure on sequences and helps identify critical evolutionary sites. The method compares K-mers between consecutive rounds, identifying instances where a K-mer from an earlier round has evolved into a different K-mer in a subsequent round defined by a specified maximum number of changes allowed per K-mer, termed ‘evolution_count’. If a K-mer in an earlier round differs from any K-mers in later by the defined ‘evolution_count’, it is considered evolved. The method generates a detailed report of the raw and evolved K-mers, their frequencies, and the positions and types of evolution.

We employed the ***Kmer_mergence*** method to aggregate reports from consecutive rounds, addressing inconsistencies in K-mer characteristics across intermediate rounds. The Kmer_evolution method, when analyzing consecutive SELEX rounds, highlights the diversity and inconsistency of intermediate-round K-mers: some appear only in the preceding round, some emerge as new in the following round, and others persist throughout the evolutionary process.

**Figure S4.**
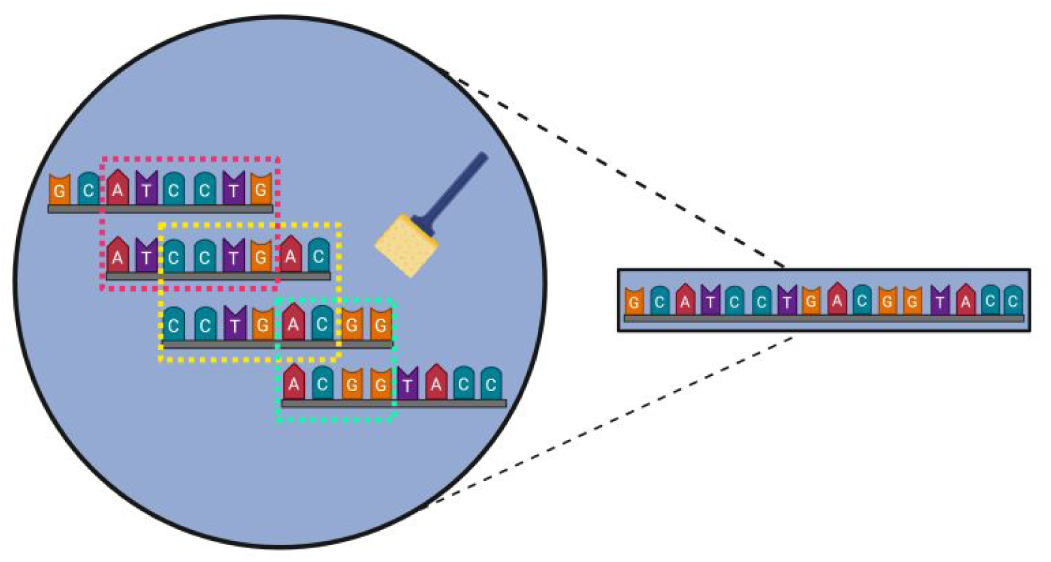
Schematic of the ***Kmer_concatenation*** method. Different colored dashed boxes in pink, yellow and green indicate the overlapping regions, which serve as the ‘junction points’ for concatenation. For example, Four short K-mers ‘GCATCCTG’, ‘ATCCTGAC’, ‘CCTGACGG’, and ‘ACGGTACC’ (left) can be automatically concatenated into one longer segment ‘GCATCCTGACGGTACC’ (right).

Figures S5 to S7 present the analysis results of thrombin aptamers selected by SELEX using the APV-Sankey toolbox. In this process, we employed a screening strategy that includes two discontinuous random sequences, with specific details available in the corresponding paper (Di Gioacchino, A., et al. Generative and interpretable machine learning for aptamer design and analysis of in vitro sequence selection. PLoS Comput Biol 2022;18(9):e1010561.). This method is significantly different from the Capture-SELEX method used for rapamycin, and this distinction is also evident in the evolutionary trends of K-mers between rounds as depicted in the result figures. Due to the shorter functional nucleic acid sequence fragments in the two discontinuous random sequences, there are fewer types of K-mers and a higher abundance of each during the evolution. This leads to differences in the statistical distribution of sequences within and between rounds. In the early rounds, before high-affinity sequences are fully selected, sequences that rank high may be eliminated in subsequent rounds. In the paper on aptamer screeing of thrombin, the affinity of the final determined aptamers was not provided, hence, Figure S6 does not provide affinity color blocks as seen in Figure 3.

**Figure S5.**
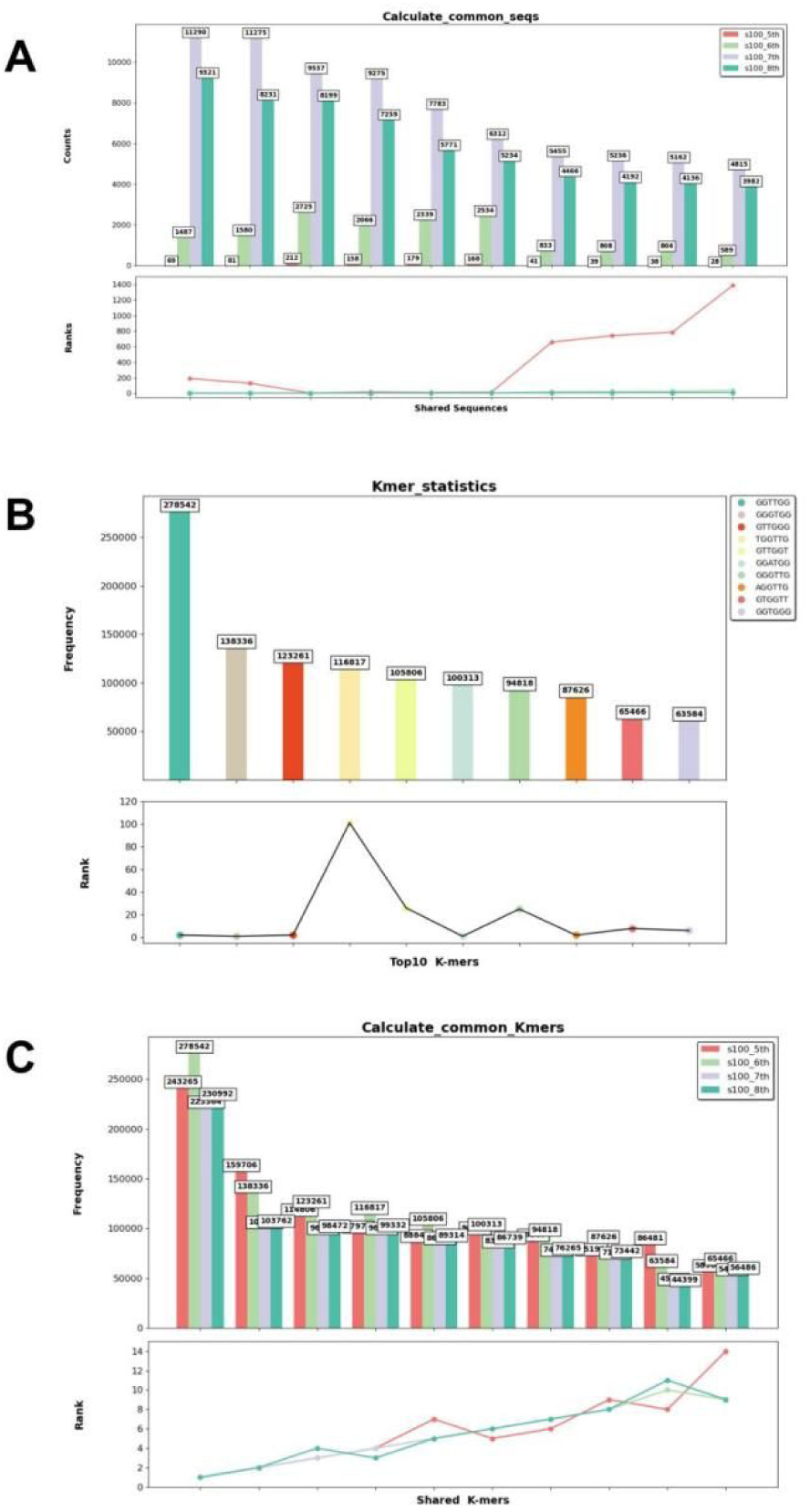
(A): The top 10 most frequent common sequences using ***Calculate_common_seqs*** method applied to the sequencing data of s100_5th, s100_6th, s100_7th, and s100_8th on aptamer screening of thrombin. The bar chart above illustrates the frequency of these common sequences in respective rounds, while the line chart below emphasizes their ranking in count for the corresponding round (the same to the following figures). (B): The top 10 most frequent K-mers using s100_8th with the ***Kmer_statistics*** method (sequence counts are not considered). (C): The top 10 most frequent common K-mers using s100_5th, s100_6th, s100_7th, and s100_8th with the ***Calculate_common_Kmers*** method (sequence counts are not considered).

**Figure S6.**
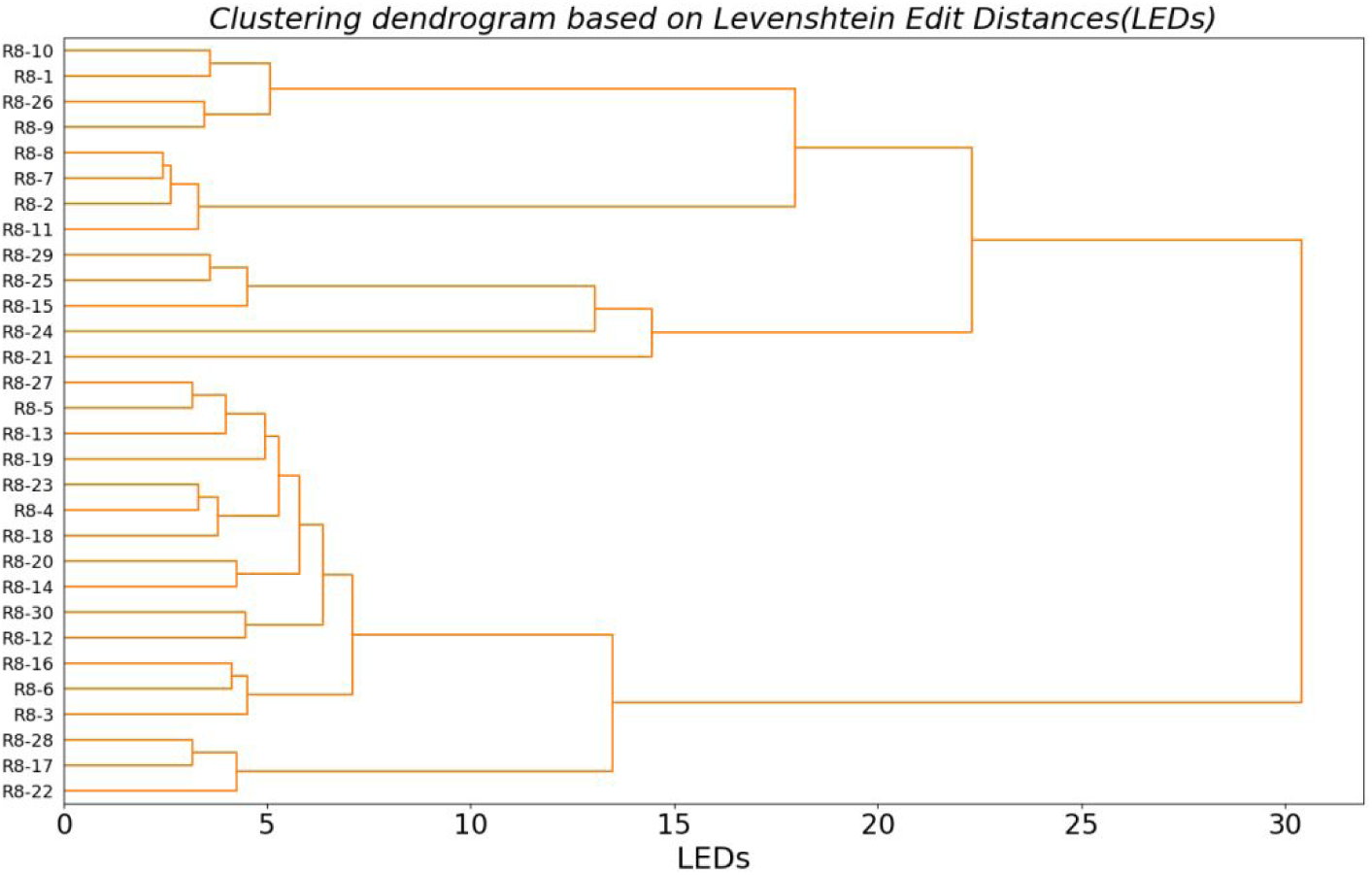
Hierarchical clustering tree generated by the ***Cluster_tree*** method to a subset of candidate sequences from s100_8th in the data of aptamer screening of thrombin.

**Figure S7.**
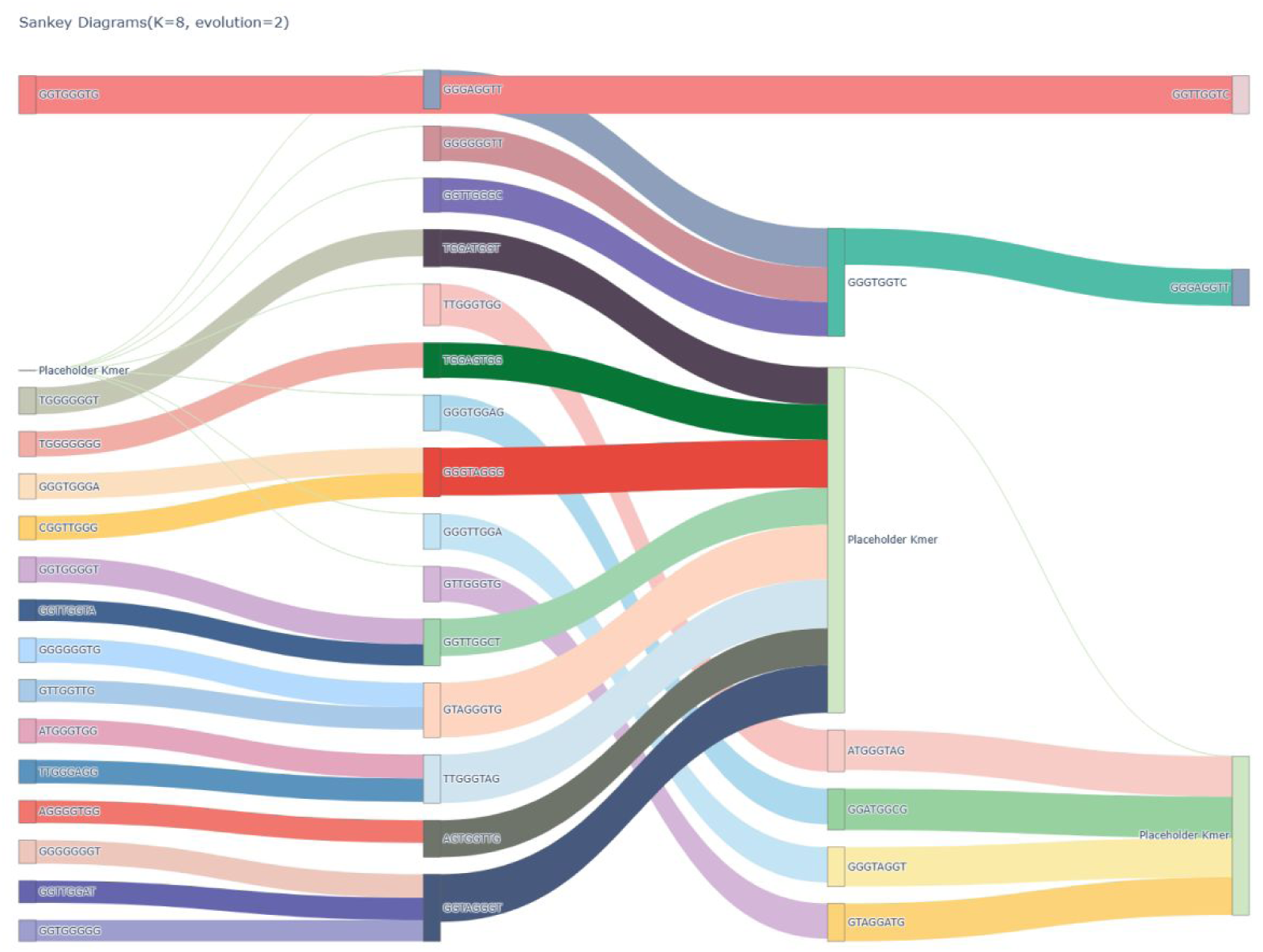
A four-layer Sankey chart constructed using s100_5th, s100_6th, s100_7th, and s100_8th in the data of aptamer screening of thrombin.

## References

1. Yu H, Zhu J, Shen G et al. Improving aptamer performance: key factors and strategies, Microchimica Acta 2023;190:255.

2. Zhou J, Rossi JJ. Cell-type-specific, Aptamer-functionalized Agents for Targeted Disease Therapy, Mol Ther Nucleic Acids 2014;3:e169.

3. Zhou J, Rossi J. Aptamers as targeted therapeutics: current potential and challenges, Nature Reviews Drug Discovery 2017;16:181–202.

4. Keefe AD, Pai S, Ellington A. Aptamers as therapeutics, Nature Reviews Drug Discovery 2010;9:537–550.

5. Ellington AD, Szostak JW. In vitro selection of RNA molecules that bind specific ligands, Nature 1990;346:818–822.

6. Tuerk C, Gold L. Systematic evolution of ligands by exponential enrichment: RNA ligands to bacteriophage T4 DNA polymerase, Science 1990;249:505–510.

7. Sharma TK, Bruno JG, Dhiman A. ABCs of DNA aptamer and related assay development, Biotechnol Adv 2017;35:275–301.

8. Lin JS, Kauff A, Diao Y et al. Creation of DNA aptamers against recombinant bone morphogenetic protein 15, Reproduction, Fertility and Development 2016;28:1164–1171.

9. Kanagawa T. Bias and artifacts in multitemplate polymerase chain reactions (PCR), J Biosci Bioeng 2003;96:317–323.

10. Luo Z, He L, Wang J et al. Developing a combined strategy for monitoring the progress of aptamer selection, Analyst 2017;142:3136–3139.

11. Shtatland T, Gill SC, Javornik BE et al. Interactions of Escherichia coli RNA with bacteriophage MS2 coat protein: genomic SELEX, Nucleic Acids Res 2000;28:E93.

12. Hoinka J, Berezhnoy A, Dao P et al. Large scale analysis of the mutational landscape in HT-SELEX improves aptamer discovery, Nucleic Acids Res 2015;43:5699–5707.

13. Chushak Y, Stone MO. In silico selection of RNA aptamers, Nucleic Acids Res 2009;37:e87.

14. Navien TN, Thevendran R, Hamdani HY et al. In silico molecular docking in DNA aptamer development, Biochimie 2021;180:54–67.

15. Lee SJ, Cho J, Lee BH et al. Design and Prediction of Aptamers Assisted by In Silico Methods, Biomedicines 2023;11:356.

16. Chen Z, Hu L, Zhang BT et al. Artificial Intelligence in Aptamer-Target Binding Prediction, Int J Mol Sci 2021;22.

17. Hofacker IL. Vienna RNA secondary structure server, Nucleic Acids Res 2003;31:3429–3431.

18. Zuker M. Mfold web server for nucleic acid folding and hybridization prediction, Nucleic Acids Res 2003;31:3406–3415.

19. Xu X, Zhao P, Chen SJ. Vfold: a web server for RNA structure and folding thermodynamics prediction, PLoS One 2014;9:e107504.

20. Biesiada M, Pachulska-Wieczorek K, Adamiak RW, Purzycka KJ. RNAComposer and RNA 3D structure prediction for nanotechnology, Methods 2016;103:120–127.

21. Wang J, Wang J, Huang Y, Xiao Y. 3dRNA v2.0: An Updated Web Server for RNA 3D Structure Prediction, Int J Mol Sci 2019;20:e4116.

22. Xu X, Chen SJ. Hierarchical Assembly of RNA Three-Dimensional Structures Based on Loop Templates, J Phys Chem B 2018;122:5327–5335.

23. Chen R, Li L, Weng Z. ZDOCK: an initial-stage protein-docking algorithm, Proteins 2003;52:80–87.

24. Morris GM, Goodsell DS, Huey R, Olson AJ. Distributed automated docking of flexible ligands to proteins: parallel applications of AutoDock 2.4, J Comput Aided Mol Des 1996;10:293–304.

25. Trott O, Olson AJ. AutoDock Vina: improving the speed and accuracy of docking with a new scoring function, efficient optimization, and multithreading, J Comput Chem 2010;31:455–461.

26. Tuszynska I, Magnus M, Jonak K et al. NPDock: a web server for protein-nucleic acid docking, Nucleic Acids Res 2015;43:W425–430.

27. Case DA, Cheatham TE, 3rd, Darden T et al. The Amber biomolecular simulation programs, J Comput Chem 2005;26:1668–1688.

28. Van Der Spoel D, Lindahl E, Hess B et al. GROMACS: fast, flexible, and free, J Comput Chem 2005;26:1701–1718.

29. Hoinka J, Berezhnoy A, Sauna ZE et al. AptaCluster - A Method to Cluster HT-SELEX Aptamer Pools and Lessons from its Application, Res Comput Mol Biol 2014;8394:115–128.

30. Alam KK, Chang JL, Burke DH. FASTAptamer: A Bioinformatic Toolkit for High-throughput Sequence Analysis of Combinatorial Selections, Mol Ther Nucleic Acids 2015;4:e230.

31. Kramer ST, Gruenke PR, Alam KK et al. FASTAptameR 2.0: A web tool for combinatorial sequence selections, Mol Ther Nucleic Acids 2022;29:862–870.

32. Dao P, Hoinka J, Takahashi M et al. AptaTRACE Elucidates RNA Sequence-Structure Motifs from Selection Trends in HT-SELEX Experiments, Cell Syst 2016;3:62–70.

33. Kato S, Ono T, Minagawa H et al. FSBC: fast string-based clustering for HT-SELEX data, BMC Bioinformatics 2020;21:263.

34. Ishida R, Adachi T, Yokota A et al. RaptRanker: in silico RNA aptamer selection from HT-SELEX experiment based on local sequence and structure information, Nucleic Acids Res 2020;48:e82.

35. Di Gioacchino A, Procyk J, Molari M et al. Generative and interpretable machine learning for aptamer design and analysis of in vitro sequence selection, PLoS Comput Biol 2022;18:e1010561.

36. Iwano N, Adachi T, Aoki K et al. Generative aptamer discovery using RaptGen, Nat Comput Sci 2022;2:378–386.

37. Jijakli K, Khraiwesh B, Fu W et al. The in vitro selection world, Methods 2016;106:3–13.

38. Kinghorn AB, Fraser LA, Lang S et al. Aptamer Bioinformatics, Int J Mol Sci 2017;18:2516.

39. Zhou Y, Qi X, Liu Y et al. DNA-Nanoscaffold-Assisted Selection of Femtomolar Bivalent Human α-Thrombin Aptamers with Potent Anticoagulant Activity, Chembiochem 2019;20:2494–2503.

40. Zhao L, Qi X, Yan X et al. Engineering Aptamer with Enhanced Affinity by Triple Helix-Based Terminal Fixation, Journal of the American Chemical Society 2019;141:17493–17497.

41. Riehmann P, Hanfler M, Fröhlich B. Interactive Sankey diagrams, IEEE Symposium on Information Visualization, 2005. INFOVIS 2005. 2005:233–240.

42. Schmidt M. The Sankey Diagram in Energy and Material Flow Management, Journal of Industrial Ecology 2008;12:82–94.

43. Yang KA, Pei R, Stojanovic MN. In vitro selection and amplification protocols for isolation of aptameric sensors for small molecules, Methods 2016;106:58–65.

44. Bruno JG. In vitro selection of DNA to chloroaromatics using magnetic microbead-based affinity separation and fluorescence detection, Biochem Biophys Res Commun 1997;234:117–120.

45. Mendonsa SD, Bowser MT. In vitro evolution of functional DNA using capillary electrophoresis, J Am Chem Soc 2004;126:20–21.

46. Park JW, Tatavarty R, Kim DW et al. Immobilization-free screening of aptamers assisted by graphene oxide, Chem Commun (Camb) 2012;48:2071–2073.

47. Nutiu R, Li Y. In vitro selection of structure-switching signaling aptamers, Angew Chem Int Ed Engl 2005;44:1061–1065.

48. Wang S, Gao H, Wei Z et al. Shortened and multivalent aptamers for ultrasensitive and rapid detection of alternariol in wheat using optical waveguide sensors, Biosens Bioelectron 2022;196:113702.

49. Sharma V, Sharma TK, Kaur I. Electrochemical detection of cortisol on graphene quantum dots modified electrodes using a rationally truncated high affinity aptamer, Applied Nanoscience 2021;11:2577–2588.

50. Ochoa S, Milam VT. Modified Nucleic Acids: Expanding the Capabilities of Functional Oligonucleotides, Molecules 2020;25:e4659.

## Reference

Di Gioacchino, A., et al. Generative and interpretable machine learning for aptamer design and analysis of in vitro sequence selection. PLoS Comput Biol 2022;18(9):e1010561.

Zhou, Y., et al. DNA-Nanoscaffold-Assisted Selection of Femtomolar Bivalent Human α-Thrombin Aptamers with Potent Anticoagulant Activity. Chembiochem 2019;20(19):2494–2503.

